# Leveraging Permutation Testing to Assess Confidence in Positive-Unlabeled Learning Applied to High-Dimensional Biological Datasets

**DOI:** 10.1101/2023.07.06.548028

**Authors:** Shiwei Xu, Margaret E. Ackerman

## Abstract

**Background:** Compared to traditional supervised machine learning approaches employing fully labeled samples, positive-unlabeled (PU) learning techniques aim to classify “unlabeled” samples based on a smaller proportion of known positive examples. This more challenging modeling goal reflects many real-world scenarios in which negative examples are not available, posing direct challenges to defining prediction accuracy robustness. While several studies have evaluated predictions learned from only definitive positive examples, few have investigated whether correct classification of a high proportion of known positives (KP) samples from among unlabeled samples can act as a surrogate to indicate a performance.

**Results:** In this study, we report a novel methodology combining multiple established PU learning-based strategies to evaluate the potential of KP samples to accurately classify unlabeled samples without using “ground truth” positive and negative labels for validation. To address model robustness, we report the first application of permutation test in PU learning. Multivariate synthetic datasets and real-world high-dimensional benchmark datasets were employed to validate the proposed pipeline with varied underlying ground truth class label compositions among the unlabeled set and different proportions of KP examples. Comparisons between model performance with actual and permutated labels could be used to distinguish reliable from unreliable models.

**Conclusions:** Like in fully supervised machine learning, permutation testing offers a means to set a baseline “no-information rate” benchmark in the context of semi-supervised PU learning inference tasks against which model performance can be compared.

## Introduction

Classification and clustering with machine learning algorithms are approaches that have been widely applied in biological and biomedical research. However, given significant experimental advances, these datasets now often present the “curse of dimensionality” [1]. While thousands of features can be easily collected in profiling studies, the number of samples is generally much smaller and often constrained by resources or biological rarity. Particularly in the context of such “wide” datasets, care must be taken to reduce potential overfitting by developing effective methods to eliminate redundant dimensionality and optimize modeling with approaches such as cross-validation and regularization. Permutation testing, in which the label or value to be modeled is scrambled among samples prior to modeling, presents an additional means to assess model robustness [2]. Permutation and modeling steps can then be repeated to generate a distribution that reflects the probability of achieving of model of a given quality by chance [3]. This distribution can be used to set a benchmark against which performance of the model trained on actual labels or values can be evaluated.

Here, we explore the use of permutation testing to address the challenging problem of establishing confidence in modeling results in the setting of semi-supervised learning (SSL) classification tasks relevant to biological and biomedical research. Specifically, we investigate a subclass of SSL tasks, known as Positive Unlabeled (PU) Learning, which has attracted the attention of researchers with datasets that consist of a small proportion of labeled positive examples and a vast majority of unlabeled samples that contain both positive and negative samples, but among which no definitive examples of true negatives are known. Summarized by *Li et al*. [4], major PU learning algorithms developed for bioinformatics tasks can be categorized into Reliable Negative Selection and Base Classifier adaptation, including but not limited to PU strategies such as bootstrapped aggregation (Bagging). Previously, we studied the empirical behavior of the transductive PU Bagging algorithm in high dimensional datasets with varied group separation and label imbalance [5]. In this context, prediction performance for PU bagging outperformed a single biased SVM classifier, which is a frequently used method in PU learning tasks, especially when the proportion of known positive (KP) samples among all samples is small [5]. Furthermore, by comparing multiple types of machine learning classifiers in PU bagging approach, we showed that the algorithm was insensitive to the choice of classifier, demonstrating an advantage when there is no ground truth label available to contribute to optimizing model selection.

However, most prior work comparing multiple PU learning-based approaches reported binary prediction performance with evaluation metrics such as accuracy, F1 score, and Matthew’s correlation coefficient (MCC), each of which requires ground truth reference labels or “True Negative” examples for validation. While these comparisons are useful, validating prediction outcomes is one of the major barriers in PU learning applications for real-world use cases, especially in biological and biomedical tasks where “true negative” is either costly or practically infeasible. Likewise, most state-of-the-art PU approaches, the ensembled model will classify the samples whether or not “true negatives” or separatable underlying clusters are present, which is considered to be one of the major factors that leads to a general failure in PU learning. In practice, there have been a limited number of studies in which researchers have attempted to validate predictions based on positive examples only (with no definitive negative examples) [6, 7]. A metric termed Explicit Precision Recall (EPR) scored the prediction by calculating the proportion of known positive samples that were predicted positive [7]. However, whether “good” prediction of positive examples can act as a reliable surrogate for “good” prediction of negative samples is certain to be context-dependent.

To this end, we propose a generalizable methodology to evaluate the robustness of positive examples to classify unlabeled samples in the absence of negative examples in the setting of transductive PU learning by integrating a previously developed PU strategy termed “Spy Positive Technique” to PU Bagging Algorithm. Proposed by Liu et al., spy positive technique was developed to determine a confident the decision boundary in PU methods that are categorized in to “Reliable Negative Selection” [4, 8] In this method, a small proportion of known positive (KP) was randomly sampled into unlabeled group and evaluate their probability to be classified as positive or negative class. Although Bekker & Davis argued that the method might not be adequate to determine decision for dataset without “sufficient” number of labeled samples, we found it suitable to be employed to evaluate the entire known positive (KP) set in PU bagging classifier by treating a small portion of KP as unlabeled and scoring them in the bagging procedure with other U set samples [9]. Moreover, we explore the use of combining this positive set evaluation and permutation tests to address the challenging problem of establishing confidence in modeling results in the setting of SSL classification tasks relevant to biological and biomedical research. Overall, in the real-world use cases, these methods were developed for understanding whether the known positive examples offer the potential to classify unlabeled instances remains a gamble without negative examples for model validation.

## Results

### Generating a distribution of positive class probability scores that reflects the null hypothesis

Inspired by the “spy” positive sample technique that was previously developed in two-step reliable negative approaches (**Fig. 1A**), we score the class 1 probability of each “spy fold” with the PU Bagging algorithm (**Fig. 1B**) [8]. The class 1 probability of each positive sample when modeled as a “spy” is further aggregated by two different scoring methods and pooled under a repeated cross-validated spy fold strategy [10]. The resulting distribution of PU Bagging scores for actual positive samples can then be compared to the distributions observed when the same inference strategy is applied after label permutation (**Fig. 1B**).

**Figure 1:**
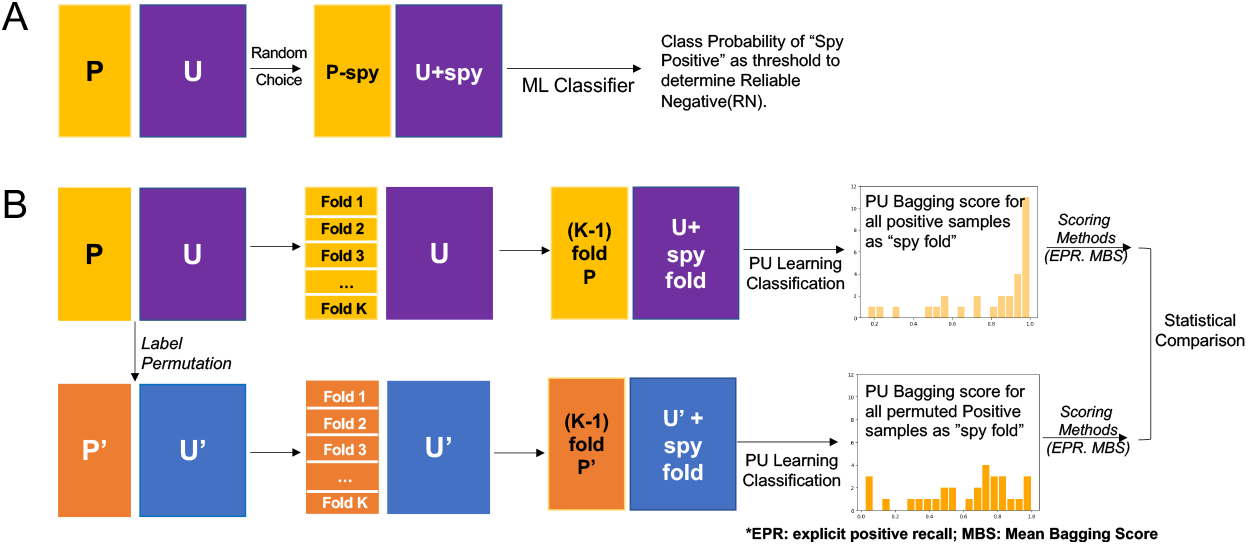
Graphical representation of the application of permutation and “spy” techniques in PU learning. **A**. Proposed by Liu et al [1], the spy method randomly samples a small percentage of the definitively labeled positive (P) class and mixes them into the unlabeled (U) set as “spies”. A classifier is then trained on the basis of the remainder of the P samples and expanded U set. The class probability of the “spy” samples is then employed to set the threshold for identification of the Reliable Negative (RN) set in a *Two-Step Reliable Negative* PU strategy. **B**. Proposed methodology to use results of permutation testing to evaluate confidence in PU learning classification. PU bagging scores across replicates and folds are calculated for “spies” and compared to scores observed for “spies” when P and U labels were randomly permuted.

### Evaluating confidence in class label inferences based on synthetic data

In previous work, we identified two characteristics that generally negatively impacted the prediction performance of PU learning methods: small separation between samples associated with ground truth label classes, and a low number of known positives among all underlying positive samples (manuscript under review). Here, we generated nine high dimensional (p≥n) synthetic datasets with different hypercube distances (high (class separation = 2), medium(class separation = 1), and a negative control (class separation = 0) and varied ground truth label class ratios (10%, 30%, 50% True Negative). The varying difficulty of these tasks based on distinctions of the underlying data captured in the first three principal components (PCs) are depicted for ground truth and positive-unlabeled classes for varying degrees of class separation, with the no class separation condition serving as a negative control (**Fig. 2A**).

**Figure 2:**
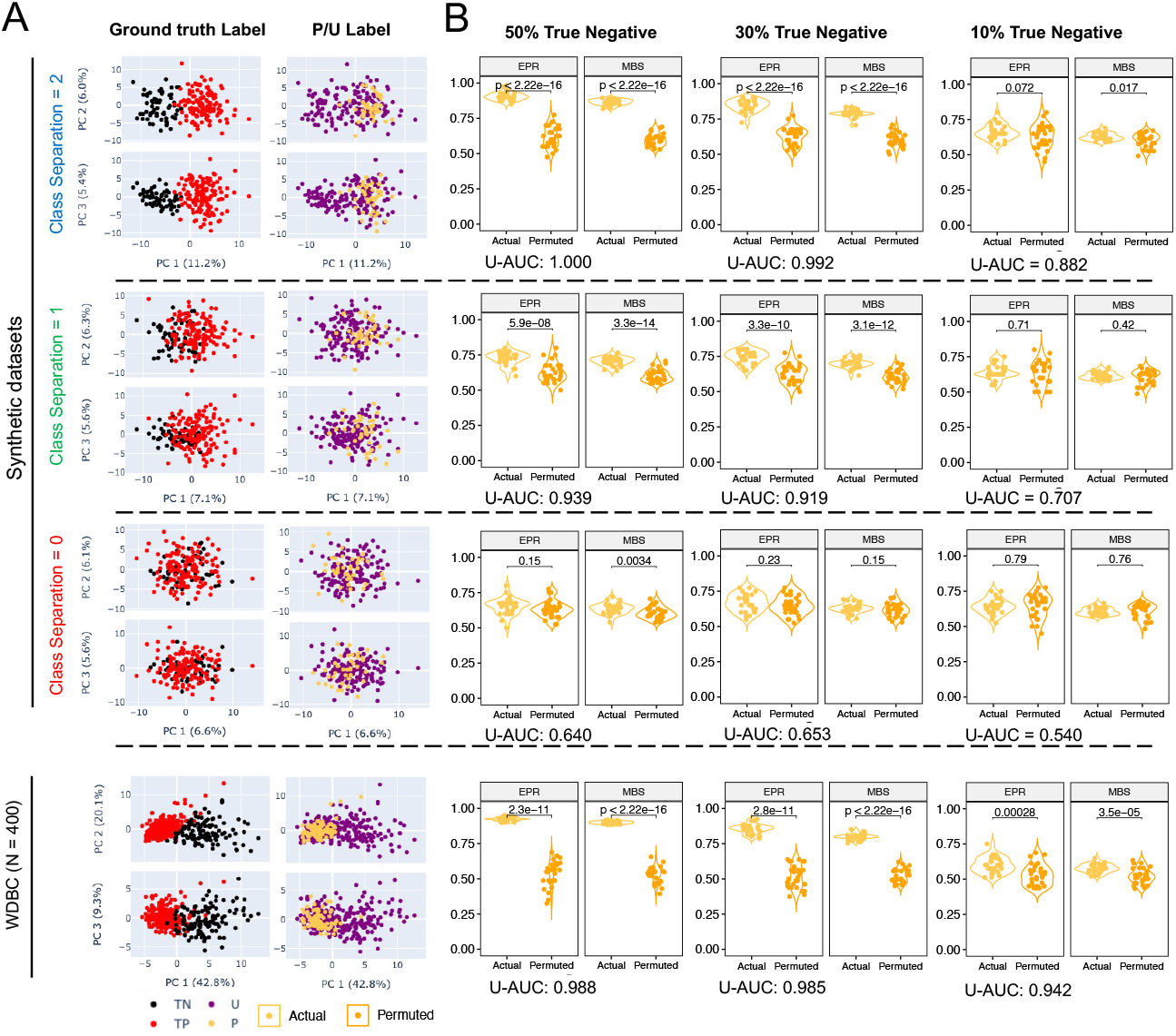
Permutation testing to evaluate model robustness across varying class separations and with extreme class imbalance. **A**. Data visualization (30% True Negative) using first three components from PCA, labeled by both ground truth class labels (left, black and red)) and P/U labels (right, purple and yellow) in the simulation settings for synthetic data with 3 different degrees of class separation (top) and Wisconsin Diagnostic Breast Cancer (WDBC) dataset (bottom). **B**. Violin plots comparing the Positive Set score under actual and permuted P/U labels for varying portions (50-10%, left to right) of True Negatives among the U set. Explicit Positive Recall (EPR) and Mean Bagging Scores (MBS) are presented and compared between actual and permuted labels (Welch’s t test, Cliff’s delta). Area under the curve for classification of unlabeled samples (U-AUC) is reported in below each test condition.

We calculated area under the receiver operator characteristic curve (ROC-AUC) between the modeled class 1 probability for known positive examples and the ground truth label for the unlabeled set in order to define expected prediction performance of inferring class assignments for the unlabeled set (**Fig. 2B inset**). As expected, the unlabeled set AUC (U-AUC) values varied from excellent (1.00) to close to random (0.64) in association with class separation when a high proportion of true negative samples were included in the unlabeled set. When the proportion of true negative samples within the unlabeled set was decreased to 10%, the U-AUC values for each condition showed a consistent reduction and exhibited the expected variation based on the level of class separation, ranging from 0.88 (excellent) to 0.54 (low to no discrimination) [11].

Modeling was then repeated after the actual PU labels were permuted multiple times. The PU bagging scores for actual and permuted known positive samples were then calculated and compared to define confidence in modeling the definitively labeled class samples (**Fig. 2B**). To thoroughly evaluate the performance the samples labeled as positive, both Explicit Positive Recall (EPR) and Mean Bagging Scores (MBS) were calculated [7]. In these simulations, we hypothesized that if the actual positive samples exhibited a distinct profile, the distribution of these scores learned for actual known positives would differ from that learned from modeling with a number-matched set of samples drawn at random. When classes were well separated (class separation = 2), distributions for actual and permuted positive samples were distinct based on tail probabilities (not shown), by Welch’s t test, Cliff’s delta estimate, which is a measure of effect size, and p-value from z-score (**Fig. 3**). As expected, performance was superior for the easier inference tasks, with high class separation and a high number (and proportion) of true negatives. Even with moderate class separation (class-separation = 1), a low proportion (10%) of true negatives yielded EPR and MBS values for actual positives that were not significantly different than those observed when labels were permuted (**Fig. 2B, Fig. 3**). Importantly, when positive and negative classes were not distinct (class separation = 0), statistically significant differences between actual and permuted EPR or MBS were generally not observed. To this end, EPR appeared somewhat less susceptible to false positive observations, suggesting that this approach is a reliable means to evaluate the level of confidence in modeling results in the absence of any true negative examples.

**Figure 3.**
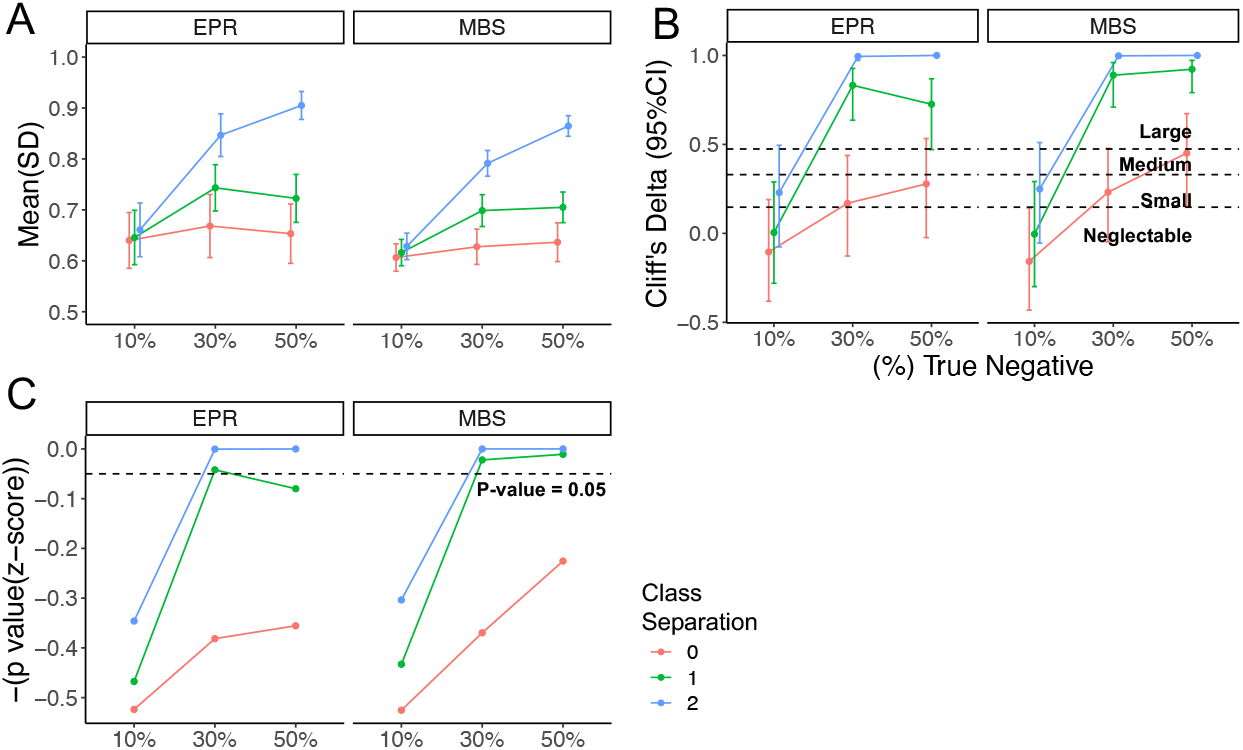
Statistical methods to evaluate scores between actual and permuted label group in the PU simulations from synthetic datasets. Explicit Positive Recall (EPR, left) and Mean Bagging Scores (MBS, right) for varying classification difficulties. Lines are colored according to class separation distance between U and P classes and x-axis indicates the composition of the unlabeled class (% True Negatives). **A**. Mean and standard deviation of the scores in actual group obtained from 30-time repetition. **B**. Cliff’s Delta estimate between scores from the actual P/U label and multiple sets of permuted labels. Error bars represented the 95% confidence interval of the estimate. Dashed line represented the boundary of *negligible / small / medium / large* difference between groups defined by Cliff’ delta statistics. **C**. Statistical significance as defined by one-tailed z test. Dashed line indicates p-value = 0.05. Lines are colored according to the underlying ground truth class separation.

### Evaluating confidence in class label inferences based on real-world data

Given results with synthetic data sets, we next applied this approach to actual biomedical data sets. Using the Wisconsin Diagnostic Breast Cancer (WDBC) dataset (**Fig. 2C**), we observed confident differences between actual known positive and permuted positive samples for all proportions of true negatives (**Fig. 2D**). In this dataset, even when EPR and MBS scores approached 0.5, the U-AUC values consistently surpassed 0.9. Additionally, statistical significance was observed across all analyzed metrics when comparing scores generated under actual and permuted P/U labels.

We further broadened our analysis to encompass additional real-world datasets (Fig. 4). In this extension, we also varied the percentage of randomly selected known positive samples ranging from as few as 10% to as many as 40% (**Fig. 4A**). This variation allowed us to examine the impact of a low proportion of known positives among all underlying positive samples and to explore the potential bias resulting from the aggregation of the bootstrapped classifier with a reduced number of training samples. To simulate the P/U learning scenario, we randomly selected a varying number of known positives from the underlying true positives in multiple real-world datasets, including the WDBC, TCGA, and Lakhashe et al. study datasets.

**Figure 4:**
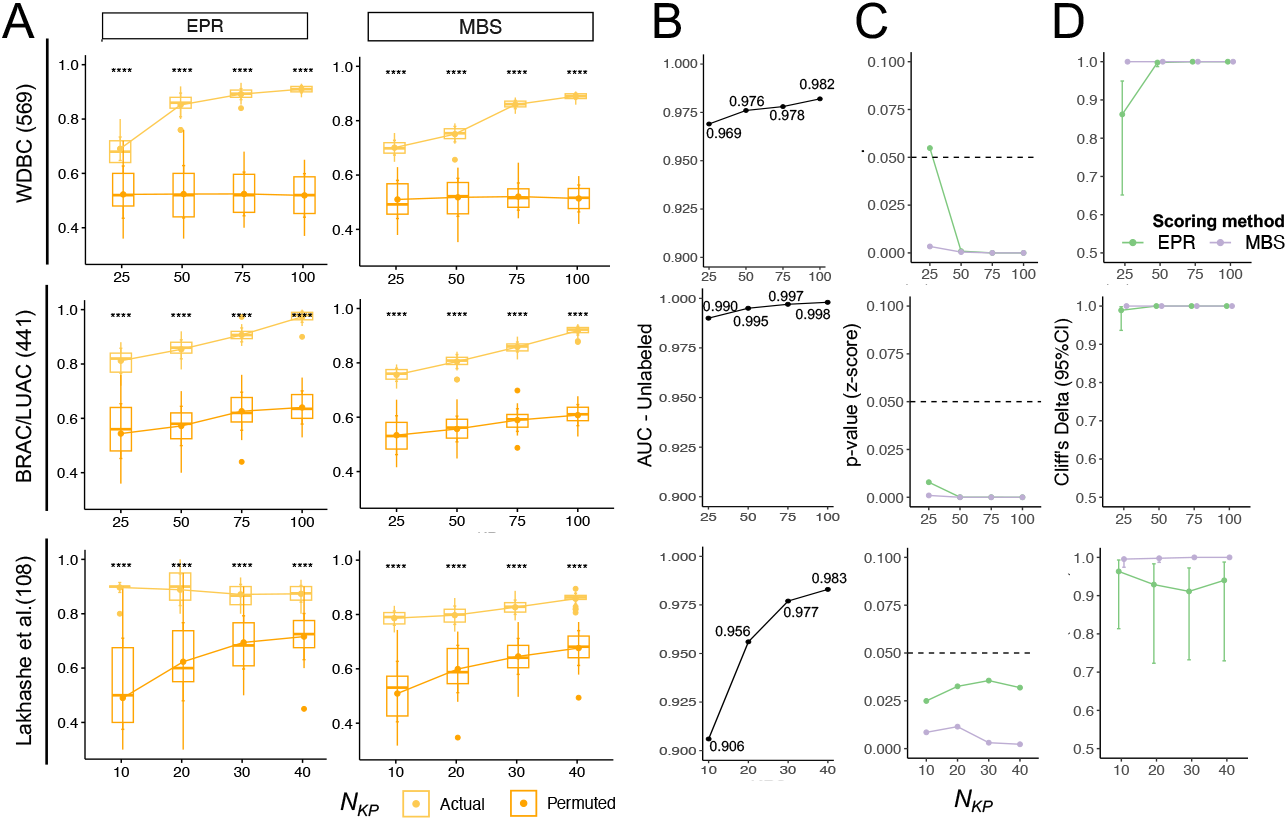
Statistical methods to evaluate scores between actual and permuted label group in real world biological datasets. **A**. Scores obtained from actual (yellow) and permuted (orange) P/U labels with two different scoring methods (EPR, left and MBS, right) under varied numbers of KP (*N*_KP_) samples for WDBC (top) BRAC/LUAC (middle) and Lakhashe et al. study dataset (bottom). Number of samples is indicated in parentheses. Boxplots depict median(bar), mean(point), interquartile range (IQR) and error bars depict mean and standard deviation (SD). Statistical significance between actual and permuted scores was defined by Welch T test (^****^: p value < 0.0001). **B**. AUC of U set samples calculated between class 1 probability using PU bagging SVM compared and ground truth label. **C**. Statistical significance from z-score between the mean score in actual label group and the distribution for scores in permuted label group. Dashed line: p value = 0.05. **D**. Effect size (Cliff’s Delta) estimate between score distributions in actual and permuted group labels.

In each of the datasets tested, high U-AUC values were observed (**Fig. 4A-B**), and actual known positive subject EPR and MBS values were statistically significantly different than observed for permuted positive subjects, regardless of the number of known positives (**Fig. 4A**). For WDBC and BRAC/LUAC, the largest data sets, increasing N_KP_ had a limited effect on permuted positive class scores, though it led to increasing values for the actual positive class label scores. In contrast, for Lakhashe et al., the smallest dataset, evaluated, increasing N_KP_ led to greater gains for permuted than actual data (**Fig. 4A**), though the distributions remained quite distinct (**Fig. 4A, C**) and a large effect size was observed (**Fig. 4D**). In sum, EPR and MBS values obtained from the actual positive class labels, even in the context of low numbers on known positive samples, were clearly distinct from results when positive labels were permuted. Concordantly, high U-AUC values, significant p-value from z-score, and large effect sizes (Cliff’s delta) provided further evidence that comparisons between actual and permuted positive class samples can provide confidence to model results in the absence of underlying ground truth label for validation (**Supplemental Table 1**).

### Permutation repetition to characterize the robustness of comparison metrics

Computational expense is a disadvantage of permutation testing in real-world applications, as it increases computational cost by a factor equal to the number of permuted label sets analyzed to confidently defined the null result distribution. To learn about the potential of wider use of the permutation test under the proposed methodology to evaluate the P set, we conducted a further investigation on the impact of the estimate from group comparison methods under an arbitrary choice of permutation repetition. Here, we specifically employed the Lakhashe et al. study dataset, a real-world humoral immune response dataset from vaccine trial with a limited sample size and wide feature space (p>n). We evaluated three different levels of permutation repetitions (30, 100, 500) and four different numbers of known positive examples to interrogate the changes in statistical significance between scores from actual labels and permuted labels (**Fig. 5A**). For all conditions, both EPR and MBS metrics indicated a large effect size (Cliff’s Delta) and high confidence (z-score or t-test). Compared to scores using the MBS value, a larger 95% confidence interval of Cliff’s Delta was observed under a lower number of permutations; however, the group difference level remained stable over the boundary of “large” (Cliff’s Delta estimate > 0.474), and a substantial change in p-value was not identified under different numbers of permutations in the p-value from the z-score, which was used to compare the mean score from the actual label and the distribution of scores from multiple permutations.

**Figure 5:**
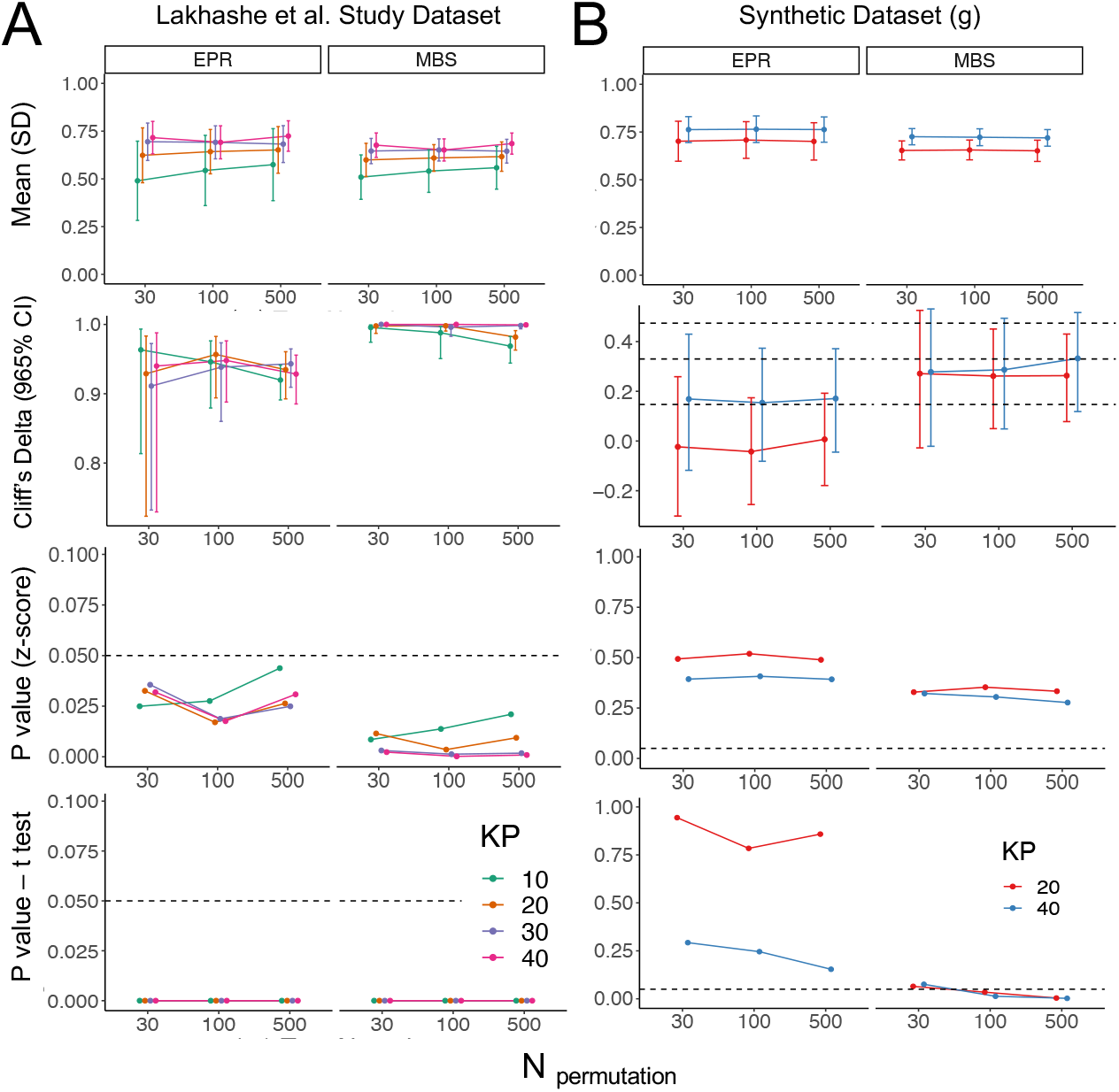
P value from the z-score remains stable regardless of the number of permutations. Statistical comparisons between scores from actual and permuted labels in Lakhashe et al. dataset and a dimension-matched synthetic dataset with varied numbers of permutation repetitions. First row: sample mean and standard deviation (error bar) of the distribution of scores under the permuted labels. Second row: Cliff’s delta estimates with a 95% confidence interval. Third row: p-value from one-sample z-score. Dashed line: p-value = 0.05. Fourth row: p-value from Welch’s T-test. Dashed line: p-value = 0.05.

In contrast to this case, in which a significant difference was obtained between scores under the actual label and permuted labels with all methods, we also investigated estimates from statistical comparison methods in a dataset where only moderate to low statistical significance was observed between scores from the actual and permuted labels. For this analysis, we generated a high dimensional synthetic dataset with a similar number of instances and attributes, but only a moderate level of hypercube distance (n:100, p:200, (%) TN:30) and poor to acceptable discrimination achieved in AUC of the unlabeled set based on the randomly selected KP from True Positive Class (TP) (KP=20: U-AUC = 0.696; KP=40: U-AUC = 0.733). With a 95% confidence interval of falling into the range from “negligible” to “medium” Cliff’s Delta, varying the number of permutations from low to high did not positively or negatively impact the estimate from this non-parametric test (**Fig. 5B**). However, a continuous decrease in p-value was observed in two-sample t-test when KP set was observed for MBS values by increasing the number of permutations, demonstrating its instability under different numbers of permutation repetitions (**Fig. 5B**). In the contrary, p-value from z-scores demonstrated stability under both larger and smaller group differences regardless of the number of permutations (**Fig. 5B**).

## Discussion

One of the primary challenges in PU learning revolves around validating binary predictions. A key point of contention is the usage of “positive” and “unlabeled” labels as substitutes for feature selection and hyperparameter tuning, especially without negative examples for validation. Another aspect of debate is the ability of PU learning-based classification methods to accurately infer the class of unlabeled samples, even when the known positive examples exhibit robust predictions within the same class. To address this dilemma, our study introduces a novel methodology that leverages and enhances traditional permutation tests within the context of PU learning. This approach provides a statistical interpretation of model confidence by comparing scores obtained from the actual P/U label with the score distribution generated from the same pipeline but with permuted P/U labels.

To thoroughly understand the empirical behavior of the proposed methodology, we generated a diversity of real-world scenarios, including underlying ground truth class separation, TP/TN label imbalance, and low percentages and numbers of KP samples to experiment with its applicability. Here, the results from both synthetic datasets and benchmark datasets demonstrated that the methodology can identify good prediction performance in the U set based on statistical significance and effect size obtained between scores from actual and permuted labels. To support its application under high dimensionality, which is a universal challenge in biological research datasets, we further investigated the impact of the statistical significance by the number of permutations, where the advantage of z-score statistics as a comparison method over two-sample independent T test, which can be arbitrarily inflated by increasing the number of permutation replicates. Z-score not only demonstrates a decreased vulnerability to manipulation based on the choice of permutation runs but also align with a more appropriate approach in principle. Comparing the score obtained from a single actual “P/U” label to multiple permuted labels, generated by randomly shuffling the label set, adheres to a more valid conceptual framework. Additionally, this study also compared two positive set scoring methods derived from the class 1 probability of the positive samples as a “spy fold”. Compared to EPR, MBS generally showed a lower standard deviation in scores from both the actual label and the permuted label, leading to improved ability to capture differences as compared to scoring with EPR. Overall, this work establishes a means to evaluate and gain or reduce confidence in PU learning inferences in the absence of known true negatives for model validation.

Nonetheless, limitations persist. Overall, low to moderate statistical significance between the mean score from the actual PU label and the distribution of the scores from permutation repetition could be generally attributed to two potential underlying issues: small separation between ground truth classes or a low proportion of true negative samples despite large separation. In the specific cases in which only 10% of true negatives existed among all the samples, we identified a rapid drop in statistical significance with multiple statistical comparison methods between positive set scores from actual P/U label and permutation repetitions, even though acceptable to excellent discrimination was suggested by the U-AUC calculated between PU bagging score of unlabeled set and ground truth. However, without a ground truth label to validate the prediction, or prior knowledge of the underlying class proportion, the methodology is not yet capable of pinpointing the exact difficulty that might inflate the risk of misclassification.

One of the potential explanations for the poor statistical significance despite separable ground truth classes and high expected prediction performance could be a joint effect from difficulty in classifying unlabeled samples with both few known positives and few true negatives, and a decreased appropriateness to employ permutation tests to evaluate a classifier under extreme imbalance. Not only a rapid drop of mean score from the actual P/U label was caused by the former, but also the distribution of the “actual” known positive and permuted known positive tended to approximate each other in the absence of very large underlying ground truth class separation, resulting in a decrease in the observed rapid drop in the statistical significance across multiple comparison methods. Consequently, the current methodology still possesses the risk of yielding “false negatives” in accurately predicting the remaining unlabeled samples using the selected PU learning methods. Thus, the current methodology still possesses the risk to give rise to “false negative” judgement on the KP set in accurately predicting the rest of the unlabeled sample using the selected PU learning methods. To address this challenge, future studies should focus on improving existing PU learning algorithms to better handle extreme underlying ground truth label imbalances. Additionally, developing enhanced scoring methods for the KP set that can effectively quantify the probability distribution may prove beneficial.

Furthermore, the methodology employed in this study was evaluated using a K-fold split approach for the positive set, with K set to 5, serving as a proof-of-concept. It is worth noting that the choice of the number of fold splits can vary depending on the specific cases. Here, we hypothesized that employing a larger “K” may lead to higher computational requirements but could result in a less biased estimation of the “score” for the positive set in transductive PU learning, considering that the number of known positives (KP) generally influences the classification performance among unlabeled samples. However, it is important to mention that we did not investigate the potential impact of “K” on the robustness reported by permutation test in the present study, leaving room for future research to supplement this aspect.

In conclusion, the effectiveness of classifying unlabeled samples using PU learning methods relies heavily on the input features provided to the PU-based classifier and the knowledge of positive examples. In this study, we introduce a permutation testing-based methodology that serves as a “gatekeeper” to assess whether the known positive samples can achieve high prediction performance in the U set. By employing a carefully selected PU-based classifier, this approach can serve as an initial step to evaluate the potential classification performance for real-world scenarios where only positive examples are available. To the best of our knowledge, our work is the first to utilize permutation tests within the PU learning framework to establish a baseline score distribution for comparison with scores obtained from the actual labels. We emphasize the versatility of our proposed method, which demonstrates its applicability across various binary class label ratios and levels of class separation. These findings provide valuable insights for the future implementation of PU learning, particularly in high-dimensional PU learning tasks.

## Methods and Materials

### Datasets for Evaluation

The datasets that were used for this study included multivariate synthetic datasets, real-world benchmark datasets from biological and biomedical studies outside the domain of vaccinology, and one real-world humoral immune response profile (**Table 1**).

**Table 1.** Description of the datasets and positive-unlabeled settings evaluated.

|  | Datasets | #Instance | # Attributes | #True Negative (%) | # Known Positive (%) |
| --- | --- | --- | --- | --- | --- |
| a | Synthetic (ClassSep = 2) | 200 | 200 | 100 (50) | 40 (20) |
|  |  |  |  | 60 (30) |  |
|  |  |  |  | 20 (10) |  |
| b | Synthetic (ClassSep = 1) | 200 | 200 | 100 (50) | 40 (20) |
|  |  |  |  | 60 (30) |  |
|  |  |  |  | 20 (10) |  |
| c | Synthetic (ClassSep = 0) | 200 | 200 | 100 (50) | 40 (20) |
|  |  |  |  | 60 (30) |  |
|  |  |  |  | 20 (10) |  |
| **d** | Wisconsin Diagnostic Breast Cancer (WDBC) | 400 | 32 | 200 (50) | 80 (20) |
|  |  |  |  | 120 (30) |  |
|  |  |  |  | 43 (10.8%) |  |
|  |  | 569 | 32 | (212) 37.3 | 25 (4.40) |
|  |  |  |  |  | 50 (8.79) |
|  |  |  |  |  | 75 (13.2) |
|  |  |  |  |  | 100 (17.6) |
| **e** | TCGA - BRCA/LUAD | 441 | 20531 | 141 (32.0) | 25 (5.67) |
|  |  |  |  |  | 50 (11.3) |
|  |  |  |  |  | 75 (17.0) |
|  |  |  |  |  | 100 (22.7) |
| **f** | Lakhashe Study Dataset | 108 | 195 | 36 (33.3) | 10 (9.30) |
|  |  |  |  |  | 20 (18.5) |
|  |  |  |  |  | 30 (27.8) |
|  |  |  |  |  | 40 (37.0) |
| **g** | Synthetic (ClassSep = 1) | 100 | 200 | 30(30.0) | 20 (20) |
|  |  |  |  |  | 40(40) |

#### Synthetic Datasets

For each synthetic dataset, profiles modeling responses for a set of 200 features for each of 200 subjects were generated to study the applicability of the proposed method in a dataset with limited sample size (p≥n) with the scikit-learn library in Python [12]. To introduce covariance, features were composed of 30% “informative” features and 70% “redundant” features, which were generated as a random linear combination of informative features. Multiple synthetic datasets were generated with varied percentages of true negative (TN) samples (10%, 30%, 50%) and varied separation (small, medium, large) between hypercubes, which were labeled either TN or true positive (TP) (**Table 1a-c**). Furthermore, a smaller dataset comprised of 100 samples with moderate hypercube separation was generated to assess the stability of statistical test results in the context of varied numbers of permutation replicates (**Table 1g**).

#### Wisconsin Diagnostic Breast Cancer (WDBC) Dataset

The WDBC dataset is comprised of 30 numeric measurements from 10 different characteristics of cell nuclei resulting from digitalized images [13]. To evaluate the proposed method in the context of varied class label ratios, three individual datasets with 400 data points from WDBC were randomly sampled from a total of 569 instances that were initially categorically labeled as benign or malignant, with 50%, 30%, and 10.8% samples as TN. For simulation purposes, samples originally labeled “Malign” were relabeled as Class 0, representing TN, and samples labeled as “Benign” were assigned Class 1, representing TP (**Table 1d**).

#### BRAC/LUAC Dataset

Datapoints labeled with tumor type *BRCA* (TP, n = 300) and *LUAD* (TN, n = 141) were selected from the PANCAN dataset, comprised of RNA Seq gene expression [14]. To speed up the training and testing time considering the high dimensionality of RNA sequencing data feature space, Principal Component Analysis (PCA) was performed to identify the top 325 principal components, which were selected to retain 95% percent of the variance in this dataset (**Table 1e**).

#### Lakhashe Study Dataset

Humoral immune profiles of responses elicited in 36 Rhesus Macaques by one of three distinct vaccine regimens across three distinct timepoints over the series of immunizations were profiled by multiplex immunoassay [15]. Among the three immunization regimens (M, K, L), samples from group L, which displayed overall vaccine efficacy, were defined as TN (n=36) (**Table 1f**).

### Positive-Unlabeled Scenario Simulation

Without external prior knowledge of sample weights or the distribution of known positive (KP) samples, Positive and Unlabeled (P/U) labels were assigned by randomly selecting KP from the TP class and leaving the rest of the data points as unlabeled (U). NumPy random seed functions in Python were used for the purpose of reproducing the results only [16].

### Analysis Pipeline

#### K-fold Spy Positive

Adapted from the concept of the “spy positive technique” in PU learning, the KP set was first randomly allocated into k folds, with one of the folds reassigned to the U set each time as a “spy fold” [17]. The PU learning method described below was then employed with the remainder of the KP samples and the updated U set (U+spy fold) to predict the Class 1 probability of all samples in the updated U set. The Bagging Scores of all “spy positive” samples were pooled. For each set of actual KP, 30 different k-fold splits were performed to define variance **(Figure 1A)**.

#### PU Learning

Transductive positive-unlabeled bootstrapped aggregation (PU bagging) described by Mordelet et al. was employed as the PU method in this study [5]. According to the number of KP in each dataset, a matched number of unlabeled subjects were randomly sampled with replacement (bootstrapped) from the U set and temporarily labeled as “negative”. A classifier was built with all KPs and bootstrapped unlabeled samples to predict the remaining Out-of-Bag (OOB) samples in the U set as negative or positive (Class 0 or 1). To improve numerical stability, prior to transformation, initial parameters for centering and scaling features were calculated from the “Bagged” samples. The PU Bagging score for each U sample was defined by aggregation from 100-time repeated bootstrapping as follows:

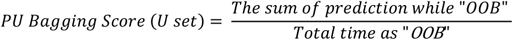

Support vector machine with the Radial Basis Function kernel (SVM-RBF) was used as a classifier in the PU bagging approach; it combines multiple polynomial kernels to project the non-linearly separatable data into higher dimension spaces to separate the targeted classes with a hyperplane created by a linear SVM. In our previous work, we showed that PU bagging with SVM-RBF classifier using the default values from Scikit-learn SVM package in python empirically achieved higher performance than linear SVM, as well as computational efficiency in high dimension synthetic and real-world biological data, without the benefit of ground truth labels in hyperparameter tuning. To indicate the expected prediction performance of the PU learning method to classify unlabeled samples, Area under the Received Operator Characteristic Curve (ROC-AUC) for the unlabeled samples was calculated between their PU Bagging scores and ground truth labels of U set samples.

#### Scoring Methods

Two different scoring methods were employed to evaluate the quality of model predictions. Defined by Cheng et al. [7], Explicit Positive Recall (EPR) is defined as the proportion of known positive examples that were predicted as “positive” among all known positive samples in the validation set. EPR was and calculated by labeling positive samples as “predicted positive” when the Bagging Score exceeded 0.5. We also consider an alternative method that lacks the requirement to set an explicit decision boundary, termed Mean Bagging Score (MBS), which is calculated as the average Class 1 probability obtained from the PU bagging classifier for positive samples.

#### Label Permutation

Permuted labels were generated by randomly shuffling the label of KP/U multiple times (repetition=30, repetition = 100, repetition = 500). For each replicate, the “permuted” positive set was assessed and scored with the procedure described above for “actual” KP samples to define the baseline performance of the proposed method for the datasets analyzed.

### Statistics

The scores obtained from the above steps were compared between “actual” and “permuted”. The distribution of scores obtained in repetition was summarized with mean and standard deviation (SD). Furthermore, both parametric and non-parametric methods were employed to compare the observed scores under the actual label to the distribution of scores under the permutated labels, including Welch’s T-test, Cliff’s delta with a 95% confidence interval, and p-value from z-score. In this study, the z-score was calculated as:

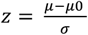

Where *μ* represents the mean score of positive sets from the 30-time repeated cross-validation process; *μ*0 represents the mean score from repeatedly permuted labels; σ is the standard deviation of scores distribution in permutation group using *n-1* degree of freedom. An upper-tailed test was performed under the hypothesis that *μ* > *μ*0.

### Visualization

PCA plots were developed and generated with “plotly” package in Python; T test annotation and plots were created with “ggpubr” package in R (version 4.2.2) [18, 19].

### Code availability

Code developed for this analysis is available on request.

## Author Contribution

<u>Investigation, coding, data visualization</u>, <u>writing-original draft</u>: S.X.; <u>Writing-reviewing and editing</u>: S.X. and M.E.A. <u>Supervision</u>: M.E.A.; <u>Conceptualization and funding</u>: M.E.A.

## Acknowledgements

This study was supported in part by NIAID R56AI165448 and P01AI162242.

