## Supplemental Material for "Leveraging Permutation Testing to Assess Confidence in Positive-Unlabeled Learning Applied to High-Dimensional Biological Datasets"

| Supplemental Tables | |
|---|---|
| Supplemental Table 1 | Statistics output from multiple comparison methods applied in Figure 2. |
| Supplemental Table 2 | Statistics output from multiple comparison methods applied in Figure 5A. |

| Dataset | Class Separation | Statistic | 50% TN | | 30%TN | | 10%TN | |
|---|---|---|---|---|---|---|---|---|
| | | | EPR | MBS | EPR | MBS | EPR | MBS |
| Synthetic datasets | 2 (easy) | Mean (SD) - Actual | 0.905 (0.027) | 0.864 (0.020) | 0.846 (0.042) | 0.791 (0.025) | 0.661 (0.053) | 0.628 (0.026) |
| | | Cliff's Delta estimate (95% CI) | 1 (0.998,1) | 1 (0.998, 1) | 0.994 (0.972, 0.999) | 0.998 (0.987, 1) | 0.229 (-0.076, 0.495) | 0.249 (-0.055, 0.51) |
| | | P value - z-score | 6.204e-05 | 3.665e-10 | 3.839e-04 | 2.839e-05 | 0.346 | 0.304 |
| | 1(medium) | Mean (SD) -Actual | 0.723 (0.047) | 0.704 (0.030) | 0.743 (0.045) | 0.699 (0.031) | 0.646 (0.053) | 0.616 (0.026) |
| | | Cliff's Delta estimate (95%CI) | 0.726 (0.471,0.869) | 0.922 (0.791, 0.972) | 0.832 (0.636, 0.927) | 0.889 (0.71, 0.96) | 0.004 (-0.281,0.289) | -0.004 (-0.299, 0.291) |
| | | P value – z-score | 0.08 | 0.011 | 0.042 | 0.022 | 0.468 | 0.433 |
| | 0 (negative control) | Mean (SD) - Actual | 0.653 (0.058) | 0.636 (0.038) | 0.668 (0.062) | 0.627 (0.035) | 0.640 (0.055) | 0.607 (0.027) |
| | | Cliff's Delta: estimate (95% CI) | 0.278 (-0.024 0.533) | 0.451 (0.155, 0.673) | 0.169 (-0.127, 0.438) | 0.231 (-0.049, 0.477) | -0.104 (-0.381 0.19) | -0.158 (-0.432, 0.143) |
| | | P value – z-score | 0.356 | 0.225 | 0.381 | 0.370 | 0.524 | 0.525 |
| WDBC datasets (400) | NA | Mean (SD) - Actual | 0.922 (0.013) | 0.9 (0.01) | 0.855 (0.034) | 0.799(0.022) | 0.6 (0.056) | 0.577 (0.031) |
| | | Cliff's Delta estimate (95% CI) | 1 (0.998,1) | 1 (0.998,1) | 1 (0.998 1) | 1 (0.998 1) | 0.546 (0.259, 0.744) | 0.6 (0.309, 0.788) |
| | | P value – z-score | 1.531e-05 | 2.613e-11 | 1.245e-05 | 5.950e-09 | 0.171 | 0.152 |

**Supplementary Table 1: Statistics output from multiple comparison methods applied in Figure 2.**

| Dataset | $N_{KP}$ | Statistics | Permutation = 30 | | Permutation = 100 | | Permutation = 500 | |
|---|---|---|---|---|---|---|---|---|
| | | | EPR | MBS | EPR | MBS | EPR | MBS |
| Lakhashe et al. study dataset | 10 | Mean (SD) - Permuted | 0.49(0.207) | 0.509(0.116) | 0.544(0.184) | 0.541(0.111) | 0.574(0.189) | 0.558(0.112) |
| | | Cliff's Delta estimate (95% CI) | 0.963 (0.814, 0.993) | 0.996 (0.974, 0.999) | 0.946 (0.879, 0.976) | 0.988 (0.951, 0.997) | 0.92(0.891, 0.941) | 0.969 (0.944, 0.983) |
| | | P value (z-score) | 0.025 | 0.009 | 0.028 | 0.014 | 0.044 | 0.021 |
| | | P value (T test) | 1.16e-11 | 4.5e-14 | 1.47e-35 | 3.27e-41 | 3.23e-135 | 1.22e-61 |
| | 20 | Mean (SD) -Permuted | 0.623(0.144) | 0.599(0.087) | 0.642(0.116) | 0.609(0.07) | 0.652(0.122) | 0.616(0.077) |
| | | Cliff's Delta estimate (95%CI) | 0.929 (0.723, 0.983) | 0.998 (0.987, 1) | 0.957 (0.894, 0.983) | 0.998 (0.99, 1) | 0.935 (0.893, 0.961) | 0.982 (0.963, 0.991) |
| | | P value (z-score) | 0.033 | 0.011 | 0.017 | 0.004 | 0.026 | 0.009 |
| | | P value (T test) | 1.92e-11 | 5.46e-14 | 2.22e-28 | 2.56e-37 | 3.46e-24 | 3.45e-30 |
| | 30 | Mean (SD) - Permuted | 0.694(0.098) | 0.646(0.066) | 0.691(0.086) | 0.651(0.058) | 0.682(0.096) | 0.645(0.062) |
| | | Cliff's Delta: estimate (95% CI) | 0.911 (0.732, 0.972) | 1(0.998, 1) | 0.938(0.86, 0.973) | 0.996(0.983, 0.999) | 0.943(0.909, 0.965) | 0.998(0.994, 1) |
| | | P value (z-score) | 0.036 | 0.003 | 0.019 | 0.001 | 0.025 | 0.002 |
| | | P value (T test) | 3.62e-11 | 1.62e-16 | 1.01e-31 | 1.36e-43 | 3.43e-31 | 8.14e-36 |
| | 40 | Mean (SD) - Permuted | 0.716(0.085) | 0.676(0.064) | 0.691(0.086) | 0.651(0.058) | 0.724(0.08) | 0.685(0.055) |
| | | Cliff's Delta estimate (95% CI) | 0.94 (0.729, 0.988) | 1 (0.998, 1) | 0.948 (0.888, 0.976) | 1 (0.999, 1) | 0.928 (0.885, 0.956) | 0.999 (0.997, 1) |
| | | P value (z-score) | 0.032 | 0.002 | 0.018 | 0.0002 | 0.031 | 0.0009 |
| | | P value (T test) | 1.85e-11 | 1.5e-16 | 1.03e-35 | 1.56e-59 | 8.25e-30 | 2.07e-45 |

**Supplementary Table 2: Statistics output from multiple comparison methods applied in Figure 5A.**
